# A microfluidic platform for sequential assembly and separation of synthetic cell models

**DOI:** 10.1101/2021.09.28.462140

**Authors:** Ran Tivony, Marcus Fletcher, Kareem Al Nahas, Ulrich F Keyser

**Affiliations:** Cavendish Laboratory, University of Cambridge, JJ Thomson Avenue, Cambridge CB3 0HE, UK

**Author notes:** These authors contributed equally to this work (Ran Tivony, Marcus Fletcher).

**Keywords:** lipid bilayer, giant unilamellar vesicles, microfluidics, artificial cell models, giant vesicle purification, bottom-up synthesis

## Abstract

Cell-sized vesicles like giant unilamellar vesicles (GUVs) are established as a promising biomimetic model for studying cellular phenomena in isolation. However, the presence of residual components and by-products, generated during vesicles preparation and manipulation, severely limits the utility of GUVs in applications like synthetic cells. Therefore, with the rapidly growing field of synthetic biology, there is an emergent demand for techniques that can continuously purify cell-like vesicles from diverse residues, while GUVs are being simultaneously synthesized and manipulated. We developed a microfluidic platform capable of purifying GUVs through stream bifurcation, where a stream of vesicles suspension is partitioned into three fractions - purified GUVs, residual components, and a washing solution. Using our purification approach, we showed that giant vesicles can be separated from various residues – that range in size and chemical composition – with a very high efficiency (*e* = 0.99), based on size and deformability of the filtered objects. In addition, by incorporating the purification module with a microfluidic-based GUV-formation method, octanol-assisted liposome assembly (OLA), we established an integrated production-purification microfluidic unit that sequentially produces, manipulates, and purifies GUVs. We demonstrate the applicability of the integrated device to synthetic biology through sequentially fusing SUVs with freshly prepared GUVs and separating the fused GUVs from extraneous SUVs and oil droplets at the same time.

## 1. Introduction

Artificial cell models like giant unilamellar vesicles (GUVs), micron-sized capsules enclosed by a phospholipid bilayer, are established as a promising tool for studying cellular phenomena and as minimal biomimetic compartments for bottom-up assembly of synthetic cells^*1-3*^. With dimensions and membrane that resembles biological cells GUVs can incorporate specific biological systems and be used as a robust biomimetic model for studying various facets of cell biology in a simplified environment under the microscope^*3, 4*^.

The relevance of GUVs to the fields of cellular and synthetic biology is perhaps best manifested by the variety of existing production and manipulation techniques^*1, 5*^. Over the last decades, different methods have been developed to allow higher degree of control over vesicle size, lipid composition, encapsulation efficiency and protein reconstitution^*5, 6*^. In ture, these techniques provide a set of tools for building diverse models for attaining unprecedented insights into membrane traits, including phase behaviour of lipids and membrane^*7, 8*^. Nevertheless, the operation of these cell-like compartments for analysing core biological processes, such as cell division, signalling and metabolism, as well as complex protein assemblies, like the nuclear pore and efflux pumps, is still very limited^*2, 3, 9*^. One of the main reasons that GUVs are yet to be broadly used for studying complex biomolecular systems, emanates from the inevitable generation of residual contaminants and by-products during the stepwise formation and manipulation of giant vesicles^*10, 11*^.

The presence of oil droplets, lipid aggregates and small unilamellar vesicles (SUVs) – typical by-products of various GUVs formation techniques^*5, 12-14*^ – introduces a large and undesirable interfacial area which is readily available to interact with the vesicles’ lipid bilayer^*15, 16*^ and interfere with the reconstitution of membrane proteins^*17*^, cytoskeletal components^*18*^, etc. Similarly, residual molecular components like surfactants, polymers, and biomolecules, can incorporate into the lipid bilayer and disrupt its integrity, organization, and properties^*19-23*^, while excess of non-entrapped fluorophores may critically interfere with data quality and reproducibility in fluorescence imaging experiments such as dye leakage measurements^*24*^. Hence, giant vesicles purification can provide a useful step for improving the efficiency and quality of measurements and successive manipulation processes.

Existing methods for purifying GUVs rely almost exclusively on conventional macro-scale approaches^*25*^ such as dialysis, differential centrifugation^*26*^, gel chromatography, membrane filtering^*27, 28*^, and Bio-Beads^*29*^. While these techniques may provide efficient separation of GUVs, they are generally inadequate for handling very small sample volumes (i.e., submicroliters), which in the context of synthetic cells construction is essential to ensure low consumption of high-value materials (e.g., extracted biomolecules, reconstituted proteins, etc.)^*30*^. Therefore, to allow efficient filtration and recovery of these mechanically sensitive vesicles, a pre-treatment stage is often implemented through dilution or solution exchange^*27*^. As such, the integration of conventional purification methods with other GUV-related processes to support a successive bottom-up synthesis of artificial cells is improbable. Nevertheless, these impediments can be overcome by using microfluidic technology.

The advantage of microfluidics lies within its ability to precisely tune and monitor miniscule volumes of fluid (order of nano to picolitres) in a micron-scale circuit of well-defined channels^*31*^. Using droplet-based microfluidic approaches, a high-throughput formation of GUVs with uniform size can be readily achieved in many types of buffers and with very high encapsulation efficiencies^*32*^. Furthermore, microfluidics offers superior handling and manipulation of GUVs through better mixing of liquids, immobilization by trapping^*33*^, and size-based sorting^*34*^. Nevertheless, on-chip filtration of giant vesicles has been mainly utilized for removing specific types of micron-size residues^*35-39*^ and, as far as we know, the simultaneous separation of various extraneous components from GUVs has yet to be demonstrated. In the context of bottom-up synthetic biology, such a purification module can be effectively implemented to improve the production of artificial cell models^*40*^, incorporation of several biomolecular systems^*3*^, and development of synthetic drug delivery systems^*41*^.

Here, we demonstrate an integrative microfluidic platform capable of continuously purifying GUVs through bifurcating a stream of vesicles mixture into three fractions: I. residual components of the GUVs suspension, II. purified GUVs, and III. washing solution (figure 1A). This fractionation approach, in essence, results in the complete replacement of GUVs solution with a contaminant free washing solution. Our approach is based on pinched-flow fractionation (PFF) adjusted for the purpose of filtering mechanically unstable cell-sized vesicles, rather than for sorting solid particles by size as it was first designed to^*34*^. To examine our device performance, we used GUVs with well-defined diameters prepared using a double-emulsion droplet-based microfluidic method, octanol-assisted liposome assembly (OLA)^*12, 42*^. Using our purification design, we were able to efficiently separate OLA-GUVs from different types of residues that range in scale and chemical properties. In addition, we managed to combine our purification module with OLA and show that, further to its ability to function as a standalone system that purifies GUVs from its suspension components, it can also be integrated with an additional microfluidic unit to establish a single device that continuously purifies freshly-prepared giant vesicles. The applicability of our integrated device to synthetic biology was demonstrated through continuously fusing SUVs with freshly prepared GUVs and subsequently separating the fused GUVs from unfused SUVs and oil droplets at the same time.

**Figure 1.**
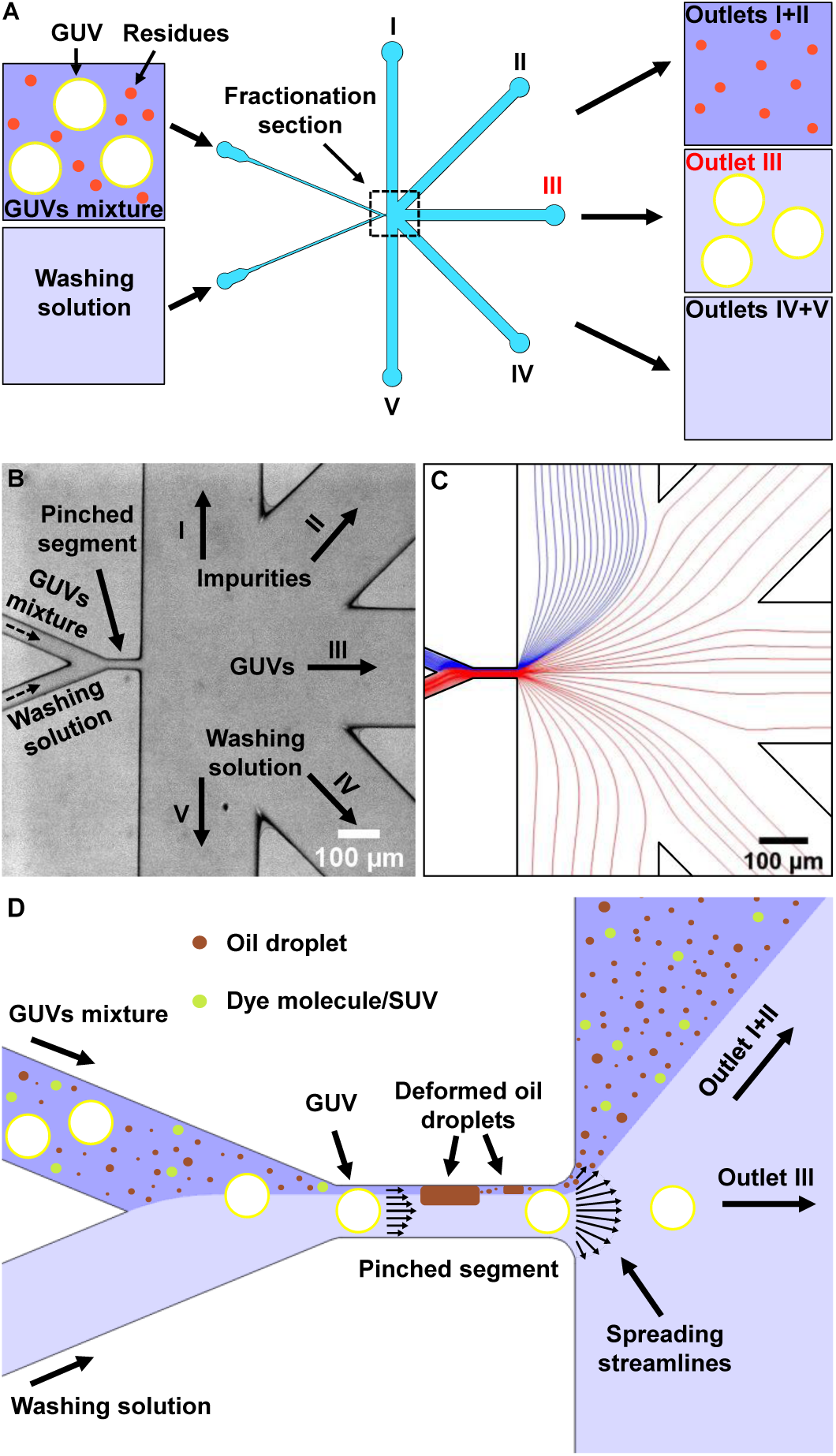
Design and basic principle of giant unilamellar vesicle (GUV) purification on-chip **A**. Schematic illustration of continuous purification of GUVs based on pinched flow fractionation (the actual design is depicted in figure S1). An isosmotic washing solution and a mixture of GUVs are perfused through the chip inlets to a fractionation section which splits them to different branch channels and outlets: impurities flow to outlets I & II; GUVs in washing solution flow to outlet III; and washing solution flows to outlets IV & V. **B**. A micrograph of the microfluidic fractionation section (marked with a dotted line square in figure 1A) with the relevant components shown in the figure, illustrating the separation concept of the purification module. **C**. Simulation of the fluid streamlines, with the relevant components depicted in the figure, showing the concept of stream bifurcation – a blue stream (i.e., GUVs mixture) is focused on the pinched segment sidewall by a red stream (i.e., washing solution). **D**. Illustration of the main concept of vesicles purification using a narrow pinched-segment with a width comparable to the diameter of GUVs. As the GUVs mixture is forced to the pinched segment sidewall by the washing solution stream, the vesicles mixture, along with all components (i.e., impurities), is separated by the spreading streamlines from vesicles whose center of mass is positioned at the microchannel centerline. In the case of large oil droplets (i.e., when *w* ≈ *a*), the viscous force inside the pinched segment stretches the oil droplets so their center of mass is shifted away from the microchannel centerline and towards its sidewall. On the other hand, the GUVs membrane is practically inextensible (hence, their surface area and volume are constant) so their center of mass is kept at (or close) the centerline.

## 2. Experimental section

### 2.1. Materials

1,2-dioleoyl-sn-glycero-3-phosphocholine (DOPC); 1,2-dioleoyl-sn-glycero-3-phospho-(1’-rac-glycerol) (sodium salt) (DOPG); 1,2-dioleoyl-sn-glycero-3-phosphoethanolamine-N-(lissamine rhodamine B sulfonyl) (ammonium salt) (18:1 Liss Rhod PE); and 1,2-dioleoyl-sn-glycero-3-phosphoethanolamine-N-(7-nitro-2-1,3-benzoxadiazol-4-yl) (ammonium salt) (18:1 NBD PE) were purchased from Avanti Polar Lipids as powder and dissolved in chloroform to a final concentration of 100mg/ml (DOPC & DOPG) and 1mg/ml (Liss Rhod PE & NBD PE). 8-Hydroxypyrene-1,3,6-trisulfonic acid trisodium salt (HPTS) was purchased from Merck and used as received. 1-octanol was purchased from sigma and used as received. Polydimethylsiloxane (PDMS) Sylgard 184 was purchased from Dow Corning and used as received.

### 2.2. Microfluidic chip design

The purification module is largely designed based on the pinched flow fractionation device reported elsewhere^*34*^ and is shown in figure 1. The microfluidic unit consists of two inlet channels that merge in a Y-junction alignment to a pinched segment, whose width is *w* = 20 μm and length is *l* = 80 μm, that connects a broadened section with five branch channels, whose width is 300 μm each (figure S1). The length of channels I, III and V is 4150 μm and of channels II and IV is 4700 μm, designed to expand the streamlines that flow to outlet III. As further elaborated in section 3.3., the integrated device consists of three sections: an octanol-assisted liposome assembly (OLA) unit for GUVs formation, a connector channel (bridge), and a purification module (figure 2A). The OLA unit was designed as specified in detail elsewhere^*43, 44*^ and contains a six-way junction which connects five channels (*w* = 20μm) – two outer aqueous channels, two lipid-octanol channels and one inner aqueous channel – to a post-junction channel (*w* = 300μm). A connector bridge channel (*w* = 500μm) is designed to combine OLA and the purification unit (same design as noted above) through their respective outlet and inlet as shown in figure 2A.

**Figure 2.**
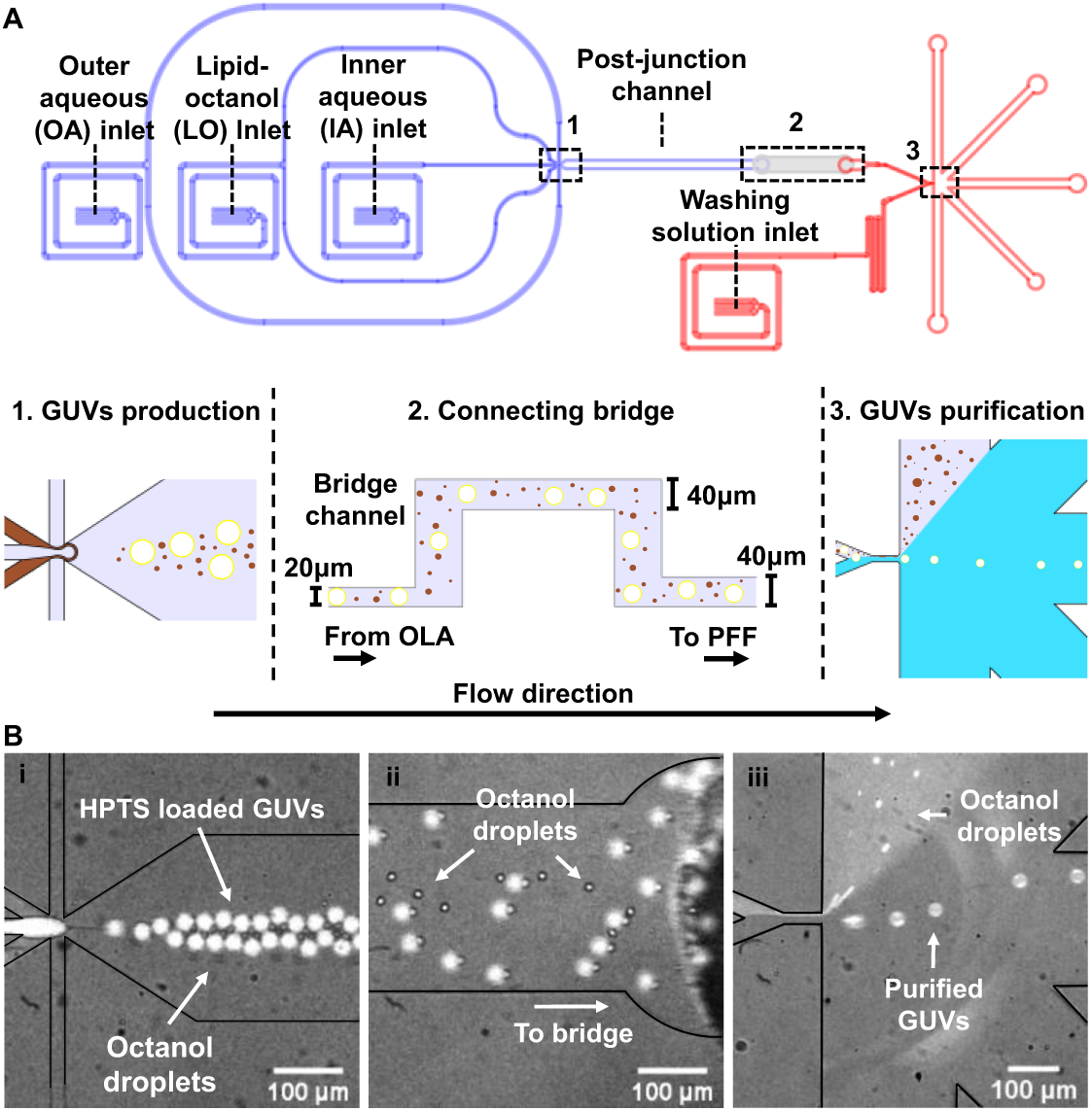
Continuous purification of GUVs using an integrated device that combines vesicle production and purification. **A**. CAD design of the integrated microfluidic device (using AutoCAD), with its principal components labelled, showing the incorporation of a GUV production module (blue; design thickness – 20μm) with the purification module (red; design thickness – 40μm), using a connecting bridge channel (light gray; design thickness – 40μm) through which the GUVs mixture flow from one unit to the other. The integrated device has 4 perfusion inlets (three for OLA – IA, LO and OA – and one for the washing solution) and 5 outlets. **B**. Fluorescent micrographs showing the sequential production and continuous purification of GUVs (labelled with HPTS (lumen) and Liss Rhod PE (membrane)) on the integrated chip. The formed HPTS-loaded GUVs (i) are drifting to the connecting bridge channel through the OLA post-junction channel outlet (ii) and reaching the purification unit where they are separated from octanol droplets using a focusing stream of a washing solution (iii). For clarity, figures (i) and (ii) show the formation and drift of vesicles in the HPTS fluorescent channel and figure (iii) illustrate the separation of these freshly-formed vesicles from oil droplets in the Lissamine Rhodamine fluorescent channel (Liss Rhod PE is in the octanol phase and in the GUVs membrane).

### 2.3. Microfluidic device fabrication

The PDMS microfluidic devices were fabricated using photolithography and soft lithography. The master mould for each design was prepared by spin-coating a thin layer of SU-8 2025 photoresist (Chestech, UK) on a 4-inch silicon wafer (University Wafer, USA). To generate a silicon master with features heights around 20 μm (OLA, production module) and 40 μm (bridge and purification module), the photoresist was spin-coated at either 2800 rpm or 1800 rpm, respectively, for 60 sec with a ramp of 100 rpm s^-1^. The wafer was then soft-baked on a hot plate at 65°C for 1 min and at 95°C for 6 min and the structures (designed in AutoCAD) were imprinted on the substrate with UV light using a table-top laser direct imaging (LDI) system (LPKF ProtoLaser LDI, Germany). After cross-linking the photoresists to form the structures the wafer was post-baked for 1 min at 65°C and for 6 min at 95°C and the structures were developed by washing away the unexposed photoresist with propylene glycol monomethyl ether acetate (PGMEA). Finally, the wafer was hard-baked for 10 min at 120°C. To generate a silicon master mould of the integrated device (figure 2A), the development process mentioned above was performed in a two-step process– the production module was prepared first by spin-coating the photoresist SU-8 2025 at 2800 rpm and after its development the purification module was printed on the same silicon wafer following spin coating at 1800 rpm. The bridge channel was prepared on a different silicon wafer by spin coating SU-8 2025 at 1800 rpm.

The PDMS chips were prepared by casting a degassed liquid PDMS (9:1 ratio with a curing agent) into the mould and then curing it for 2 hrs at 60°C. Perfusion inlets and outlets holes in the PDMS chip were created using 0.75 mm and 1.5 mm biopsy punches (WPI, UK), respectively. Finally, the chip was bonded to a PDMS-coated cover slip following their exposure to oxygen plasma for 10sec (100 W plasma power, 25 sccm; plasma etcher, Diener Electric GmbH & Co, KG). The integrated PDMS chip was prepared as described above and the bridge channel (PDMS) was subsequently bonded following their exposure to oxygen plasma for 1 min.

### 2.4. Chip operation and data acquisition

The microfluidic chips were operated on an Olympus IX73 inverted microscope and fluids in the chip were controlled via a pressure-driven pump (MFCS-EZ, Fluigent GmbH, Germany) using either 2 pressure ports (standalone purification device, figure S1) or 4 pressure ports (integrated device, figure 2A), where flow rates of perfused fluids were tuned and monitored in real-time using an accompanying MAESFLOW software. The perfusion of fluids from their reservoirs (Micrewtube 0.5ml, Simport) and through the microfluidic chip was done via a polymer tubing (Tygon microbore tubing 0.020” ID x 0.060” OD, Cole-Parmer, UK) and a metal connector tip (removed from a dispensing tip; Gauge 23 blunt end, Intertronics). Before running the chip, the complete wetting of all branch channels was confirmed, and trapped air was removed if required, to ensure the correct spreading of fluid streamlines in the broadened section. In a typical chip operation, 200 μl of GUVs mixture (when using the standalone device shown in figure S1) and 1.5 ml washing solution (inner aqueous solution; see next section) were added to their reservoirs and perfused through the chip.

Images and videos were acquired by a Photometric Evolve 512 camera controlled via an open-source software μManager 1.4, using a 10x air objective (Olympus UPLFLN). For tracking HPTS and Liss Rhod PE fluorescence, FITC and Texas red filter cubes (Chroma) were used, respectively, with a wLS LED lamp (Q-Imaging) as the light source unit. An Olympus FluoView FV1000 Confocal Laser Scanning Microscope was used to image giant vesicles, fluorescently labelled with NBD PE or Liss Rhos PE lipids, through excitation with a laser at 488nm and 559 nm, respectively. All images were analyzed using ImageJ.

### 2.5. Octanol-assisted Liposome Assembly (OLA)

High-throughput preparation of monodispersed GUVs with well-defined diameters was achieved using octanol-assisted liposome assembly (OLA), as explained in detail elsewhere^*12*^. Briefly, the formation of vesicles on-chip was controlled via a pressure-driven pump by which flow rates of all three phases (inner aqueous (IA), lipid-octanol oil (LO), outer aqueous (OA)) could be tuned and monitored in real-time. The corresponding chip inlets and design are shown in figure 2A.

For all experiments, the base solution used for the IA and OA phases (prepared in Milli-Q water) consisted of 100mM HEPES, 200mM Sucrose and 15% v/v glycerol, titrated with 1M NaOH to reach a final pH value of 7.8. The OA phase (pH = 7.8) also contained 50mg/mL of Kolliphor P-188 (Sigma-Aldrich, UK). For each OLA experiment, a total volume of 200 μl was used for the IA and OA phases and 100 μl for the LO phase. To fluorescently label the GUVs lumen we encapsulated 10μM HPTS (in IA), a membrane impermeable pH sensitive dye. The LO phase comprised of 4μL of a lipid stock solution (100mg/ml DOPC:DOPG in ethanol; 3:1 v/v ratio), 1μL of a fluorescent-lipid solution (1mg/ml of either Liss Rhod PE or NBD PE in chloroform), and 95μL of 1-octanol.

### 2.6. Vesicles electroformation

Giant unilamellar vesicles (GUVs) were prepared by electroformation using a Nanion Vesicle Prep Pro setup. 1,2-diphytanoyl-sn-glycero-3-phosphocholinelipid (DPhPC) with 1,2-dipalmitoyl-sn-glycero-3-phosphoethanolamine-N-(lissamine rhodamine B sulfonyl) (16:0 Liss Rhod PE) from Avanti Polar Lipids were dissolved in chloroform DPhPC/Liss Rhod PE (in a ratio of 265:1). 60 μl of the lipid mixture at 5mg/ml was spincoated on the conducting surface of an Indium Tin Oxide (ITO) coated glass slide (Nanion/Visiontek). The chloroform was evaporated for 1 hour in a desiccator and then 600μl of the sucrose solution (100 mM sucrose in MilliQ water) was deposited within the O-ring chamber which is sealed with another ITO coated slide. The electroformation chamber was then connected to the Nanion Vesicle Prep Pro and the electroformation protocol was carried out at 37°C and proceeds in 3 steps: (i) The a/c voltage increases linearly from 0 to 3.2 V (p–p) at 10 Hz over 1 hr. (ii) The voltage stays at 3.2 V (p–p) and 10 Hz for 50 min. (iii) The frequency decreased linearly to 4 Hz over 10 min and is maintained for another 20 min.

### 2.7. Chip simulation

The steady state Navier-Stokes equations were solved using the finite element modelling software COMSOL multiphysics (version 4.4). The ‘In-built’ physics modelling module, ‘Laminar Flow’ was used with the 2D microfluidic chip CAD design defining the domain for the solution. Two inlet velocities were defined as boundary conditions for both the washing solution channel (*v*_*ws*_) and GUVs mixture channel (*v*_*GUUVs*_). The five open outlets (I-V) of branch channels were held at a constant atmospheric pressure boundary condition, *p*_*out*_ = 1 bar. To simulate the streamlines in the purification chip, *v*_*GUVs*_ was held constant and *v*_*ws*_ adjusted through a range of values.

### 2.8. FRET measurements

Negatively charged GUVs (DOPC:DOPG, 3:1) with 1mol% DOPE-NBD and different concentrations of DOPE-Rh, varying between 0 to 0.5 mol%, were prepared using OLA. The sucrose containing vesicles were then settled to the bottom of an incubation chamber in an isosmotic glucose solution. The relative FRET efficiency *E*_*FRET*_ was extracted from measuring the fluorescence intensities of DOPE-NBD and DOPE-Rh following the excitation of NBD with a 488nm laser, using an Olympus FluoView FV1000 Confocal Laser Scanning Microscope. Since each experiment generated a set of images containing 50-100 GUVs. Therefore, to extract distributions of relative FRET efficiency, we wrote a custom python script that automatically detects GUVs through the Hough Circle Transform algorithm and measure the mean intensity of NBD and Rh from the equator of vesicles.

## 3. Results and discussion

### 3.1. Principle of GUVs purification on-chip

We developed a microfluidic module that employs pinched flow fractionation (PFF) for the continuous purification of giant unilamellar vesicles (GUVs). As schematically illustrated in figure 1A, our approach is based on the precise separation of a GUVs mixture into three different fractions – residual components, purified GUVs, and an isosmotic washing solution. The actual layout of our design consists of three parts: two microchannels, arranged in the form of a Y-junction, through which the GUVs mixture (with its extraneous substances) and an isosmotic washing solution are perfused and merged; a pinched segment as the main filtering element; and a broadened section of five channels that spatially divide the obtained fractions into different outlets (figure 1B and figure S1). Typically, PFF has been exploited for sorting or separating different types of particulates based on their size^*37, 45*^. Through focusing a flow of particles on one sidewall of a wider pinched segment, particles with various diameters can be aligned in a slightly different lateral position within the microchannel. Consequently, the focused particles can be separated according to their sizes, without clogging the microchannel, by flowing along the streamline that passes through their center of mass^*34*^.

We exploit the same fundamental principles of PFF and laminar flow which, in essence, enable to bifurcate a stream of fluid (the GUVs mixture in our case) into two separate currents as follows. Figure 1C illustrates this concept by showing the theoretical profile of streamlines in our system following the perfusion of two streams, blue and red, each at a different flow rate. By using the red flow to focus the blue flow on the pinched segment sidewall, the blue fluid can be specifically directed to outlet I while the red fluid occupies the rest of the broadened section volume. To adjust our device for the purpose of separating GUVs from their mixture, we narrowed the pinched segment so its width will be comparable to the diameter of vesicles we seek to purify (figure 1D). As a result, GUVs that enter the pinched segment will have their center of mass positioned very close or at the centerline of the microchannel. Hence, once arriving to the broadened section, they will drift along the streamlines that flow to outlet III. On the other hand, the pinched mixture with all its other components (dissolved fluorescent molecules, SUVs/LUVs, oil droplets, etc.) will follow along the spreading streamlines that flow to outlet I or outlets I & II, as noted above. This approach capitalizes on the unique properties of lipid vesicles which, unlike rigid particles, can bend to fit into a narrower channel and glide across it while sustaining the resultant viscous stresses, as elaborated below. Consequently, and as will be shown later, a range of vesicle diameters can be purified from different types and sizes of impurities using a single pinched segment, and without clogging it.

### 3.2. Continuous production and purification of GUVs on-chip

To examine the performance of our purification device, we used octanol-assisted liposome assembly (OLA), a microfluidic-based platform for preparing GUVs^*12*^ (also see experimental section). While this promising technique offers several important advantages – rapid and controlled production of GUVs with well-defined diameters, an excellent encapsulation efficiency, and the use of a biocompatible organic solvent – it also generates a large amount of octanol droplets as a by-product of the vesicle preparation process. These oil droplets can deteriorate the stability of GUVs, clog the microfluidic channels, and increase the background fluorescence noise^*35*^. In addition, the presence of nanometric to micron sized lipid-carrying oil droplets introduces a large (undesirable) phospholipid monolayer surface area which can potentially interact with fluorescent molecules, membrane proteins and other biologically relevant components. Therefore, it is essential to filter out all octanol droplets (and dissolved octanol) following the production of OLA-GUVs.

Very recently, a density-based microfluidic filtering technique was designed to separate GUVs from oil droplets that formed as a by-product during vesicle formation on-chip by octanol-assisted liposome assembly (OLA)^*35*^. In principle, this purification technique applies small negative pressures to pull GUVs from the bottom of an outlet reservoir while leaving the oil droplets to float upwards. However, since this approach strictly relies on density differences between vesicles and oil droplets, it was found to be inappropriate for filtering out small oil droplets (<4μm) and dissolved octanol molecules^*35*^. Hence, such a density-based approach is also unsuitable for excluding solutes like surfactants, proteins and fluorescent dyes or materials such as SUVs/LUVs and micelles that have densities comparable to GUVs.

We developed an integrated microfluidic device that enables to continuously prepare and purify monodispersed GUVs (*R* = 10.3 ± 1.1 μm, see figure 5B) with high-throughput by combining the purification module with OLA (figure 2). To connect the two sections, we bonded a connecting microchannel (bridge) to the integrated chip prior to GUVs production using OLA (see experimental section). The use of a bridge allows us to divide the two microfluidic units while pre-treating the OLA post-junction channel with polyvinyl alcohol (PVA)^*42*^ – an essential step that prevents the clogging of the post-junction channel by adhesion of octanol droplets to the PDMS walls^*35*^. Without initially dividing the two microfluidic parts, PVA accumulates in the purification chamber, rendering it non-functional and thus preventing the successful integration of OLA with other PFF-based separation techniques^*35*^. As illustrated in figure 2A (bottom panel), once vesicle production begins, freshly formed GUVs and octanol droplets flow through the connecting bridge and reach the purification chamber that continuously separate them by using an isosmotic focusing stream of washing solution (inner aqueous solution) to minimize the development of osmotic imbalance. Since the flow rate in the GUVs mixture channel *Q*_*GUVs*_ is coupled to the production rate of vesicles in the OLA section, we measured it during a typical GUVs production and found that *Q*_*GUVs*_ = 33.4 ± 3.7 *μl hr*^−1^ (SI and figure S2).

Figure 2B demonstrates the continuous purification of freshly prepared DOPC:DOPG (3:1 molar ratio) GUVs from octanol droplets, where the pinched segment has a width of w = 20μm and a height of h = 40μm (see also video S1). As can be seen in figure 2B (iii), once the OLA mixture passes through the pinched segment it splits into two separate streams – one consists of only GUVs while the other contains octanol droplets, dissolved octanol molecules (water solubility of 1-octanol is 0.46g/L), and dissolved poloxamer P188 (see experimental section). Remarkably, even though PFF-based separation techniques are intrinsically designed to separate objects by their size^*45*^, we found that oil droplets with diameters larger than the pinched segment’s width can be filtered out, while GUVs of similar size drift along the streamlines to outlet III (figure S3). This observation can be explained by the different behaviour of oil droplets and vesicles under shear stress in a Poiseuille flow (i.e., pressure-driven laminar flow). In a rectangular microchannel, hydrodynamic stresses act to elongate oil droplets mainly along the flow direction whereas the droplet’s constant interfacial tension *σ* creates a strong lateral interfacial force that resists its extension along its width and height^*46, 47*^. The elongation of droplets is further amplified when the ratio between the viscosity of the droplet phase *η*_*d*_ and the viscosity of the continuous phase *η*_*c*_ is larger than unity^*47-49*^, as in the case of octanol and water (*λ = η*_*d*_*/η*_*c*_ ≈ 7.5)^*50*^, and when the drop is stabilized by surfactants that act to reduce the interfacial tension^*51*^ (SI). Consequently, their center of mass can, in principle, be shifted by the focusing stream away from the microchannel centerline and to outlets I and II, as schematically shown in figure 1D. On the other hand, vesicles are enclosed by an inextensible fluid membrane, which is typically stretched unless exposed to hypertonic environment (normally, this is not the case with freshly prepared vesicles), so their surface area and spherical volume are kept nearly constant in a Poiseuille flow. Consequently, their reduced volume (a measure of vesicle asphericity) largely remains close to unity *𝒱* = *V*_*GUV*_*/V*_*sphere*_ *≅ 1* when crossing the pinched segment (figure 2B (iii) shows that the vesicles retain their spherical shape after the pinched segment), where *V*_*GUV*_ and *V*_*sphere*_ are the vesicle volume and the volume of a sphere with the same radius^*52, 53*^. Altogether, the large membrane stretching modulus^*54, 55*^ and incompressibility of vesicles (as *𝒱 ≅ 1*), suggest that GUVs may only slightly deform in the pinched segment (SI) while keeping their center of mass close to the microchannel centerline.

Overall, the separation of GUVs from larger oil droplets along with their different response to shear stress, implies that our purification approach is not entirely based on size differences between vesicles and impurities but also on their deformability. The greater number of separation characteristics, broadens the utility of the purification module and allows the nearly complete separation of GUVs from the oil phase which includes dissolved oil molecules and droplets with diameters ranging from nanometers to several tens of microns.

### 3.3. Efficiency of GUVs purification

To examine the efficiency of the continuous purification process we encapsulated a water-soluble membrane-impermeable pyranine dye (HPTS) in the GUVs lumen and subsequently filtered the fluorescent vesicles from free (non-entrapped) HPTS, as shown in figure 3A (see also video S2). We note that even though OLA has an excellent encapsulation efficiency, free HPTS is still released to the external solution mainly due to vesicle rupture events and flow instabilities in the six-way junction during vesicles production. We quantified the purification efficiency by comparing the fluorescence intensity of the external solution in outlets I-III. As can be seen in figure 3B, while large intensities were measured for outlets I (*f*(I) = 3250 ± 415) and II (*f*(II) = 2167 ± 335), a very low background intensity was measured in the external solution of outlet III (*f*(III) = 48 ± 88). Although the very low background fluorescence intensity in outlet I may suggest that GUVs separation from HPTS was not complete, it is most likely that also other factors, such as out-of-focus GUVs and HPTS discharge due to bursting of filtered vesicles, contributed to the apparent signal. Still, by comparing the fluorescence intensities obtained from outlet III and I (outlet I reflects the true concentration of free HPTS in the GUVs mixture), we found that GUVs can be continuously purified from HPTS with an excellent separation efficiency of *e* = 1 − f(*III*)⁄*f(I)* = 0.99. The efficient exclusion of a solute like HPTS demonstrates the nearly complete exchange of vesicles external solution, indicating that purification is not limited to specific types or sizes of residues but can be applied to remove other types of solutes and dispersed components with similar efficiencies.

**Figure 3.**
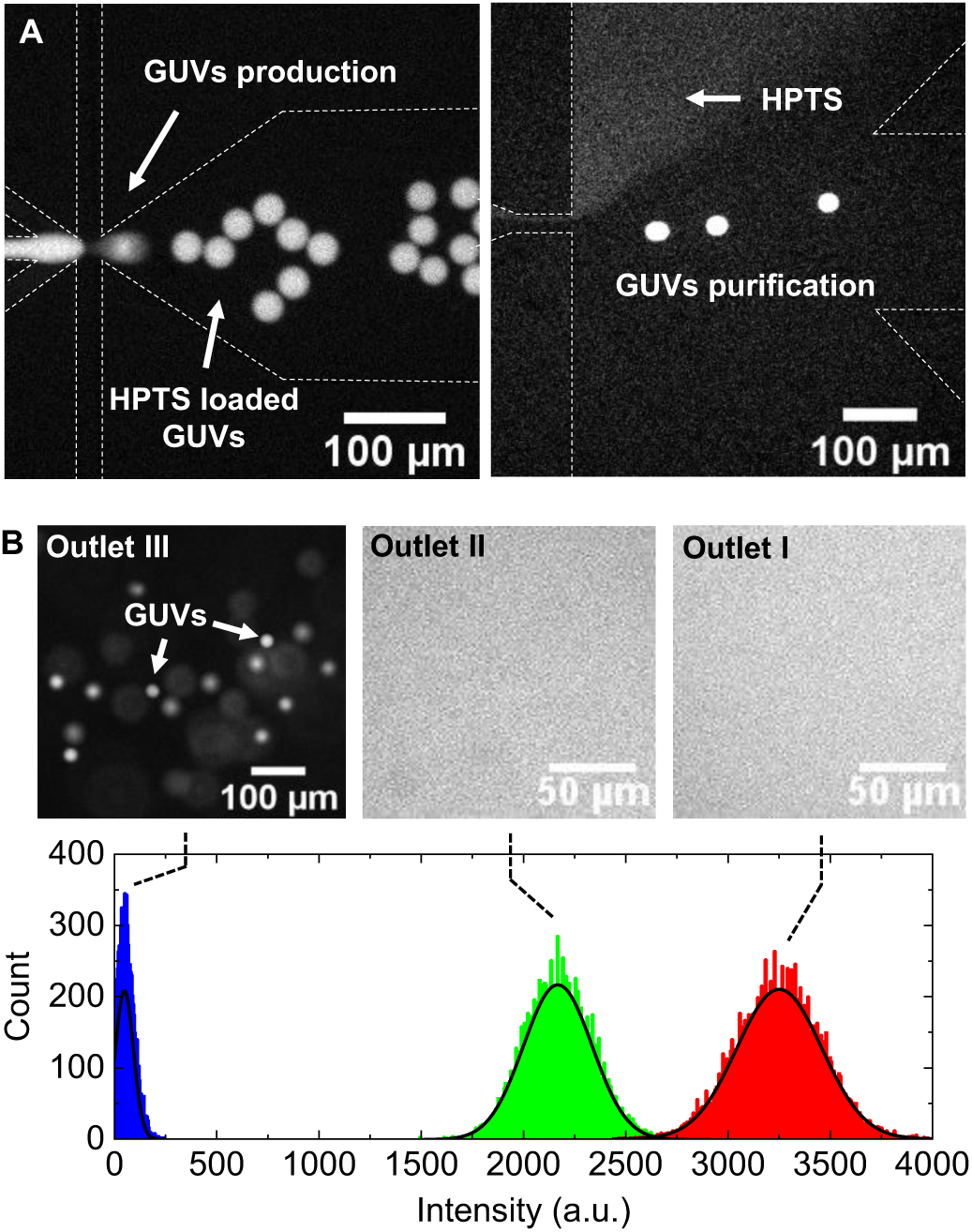
Purification efficiency and vesicles recovery. **A**. Fluorescent micrographs depicting the sequential production (left image) and purification of HPTS-loaded GUVs from free HPTS (right image) on an integrated device. **B**. The upper panel shows fluorescent micrographs of the fraction collected from each outlet (I, II, & III). The bottom panel indicates the pixel count for the fluorescence intensity in each outlet, where the black solid lines are the best fit of a gaussian distribution to the data. In outlet III, the fluorescence intensity was obtained by measuring the background signal, excluding focused and blurred (out-of-focus) GUVs, and for outlets I & II the intensity was measured over a similar pixel area as in outlet III.

### 3.4. Mechanical stability of GUVs during purification

Flow velocity is an important parameter that may critically affect the separation efficiency and recovery of GUVs during the purification process. For instance, the flow velocity ratio between the two opposing streams in the Y-junction (figure 1D) dictates the thickness of the pinched fluid and consequently the bifurcation angle *θ* (inset to figure 4A) which signifies the magnitude of spatial separation between vesicles and their mixture. While large values of *θ* imply an increased spatial separation, very high flow velocities in the pinched segment may also impose high shear stresses and, potentially, bursting of vesicles. To assess the optimal range at which GUVs can be purified while remaining intact, we first measured the bifurcation angle at different flow rate ratios between a washing solution and a fluorescent (HPTS) solution. As depicted in figure 4A, the bifurcation angle increases as the flow rate ratio between the washing solution and GUVs mixture streams *(Q*_*ws*_*/Q*_*GUVs*_) rises, where each value of *θ* indicates the specific outlets to which the HPTS solution (or impurities) flows. For instance, the black arrow in figure 4A indicates a flow rate ratio of *Q*_*ws*_⁄*Q*_*GUVs*_ ≈ *1*.*3* at which the HPTS solution flows to outlets I and II. Similarly, at ratios larger than *∼2*.*4*, as indicated by the red arrow, the HPTS stream flows only to outlet I.

**Figure 4.**
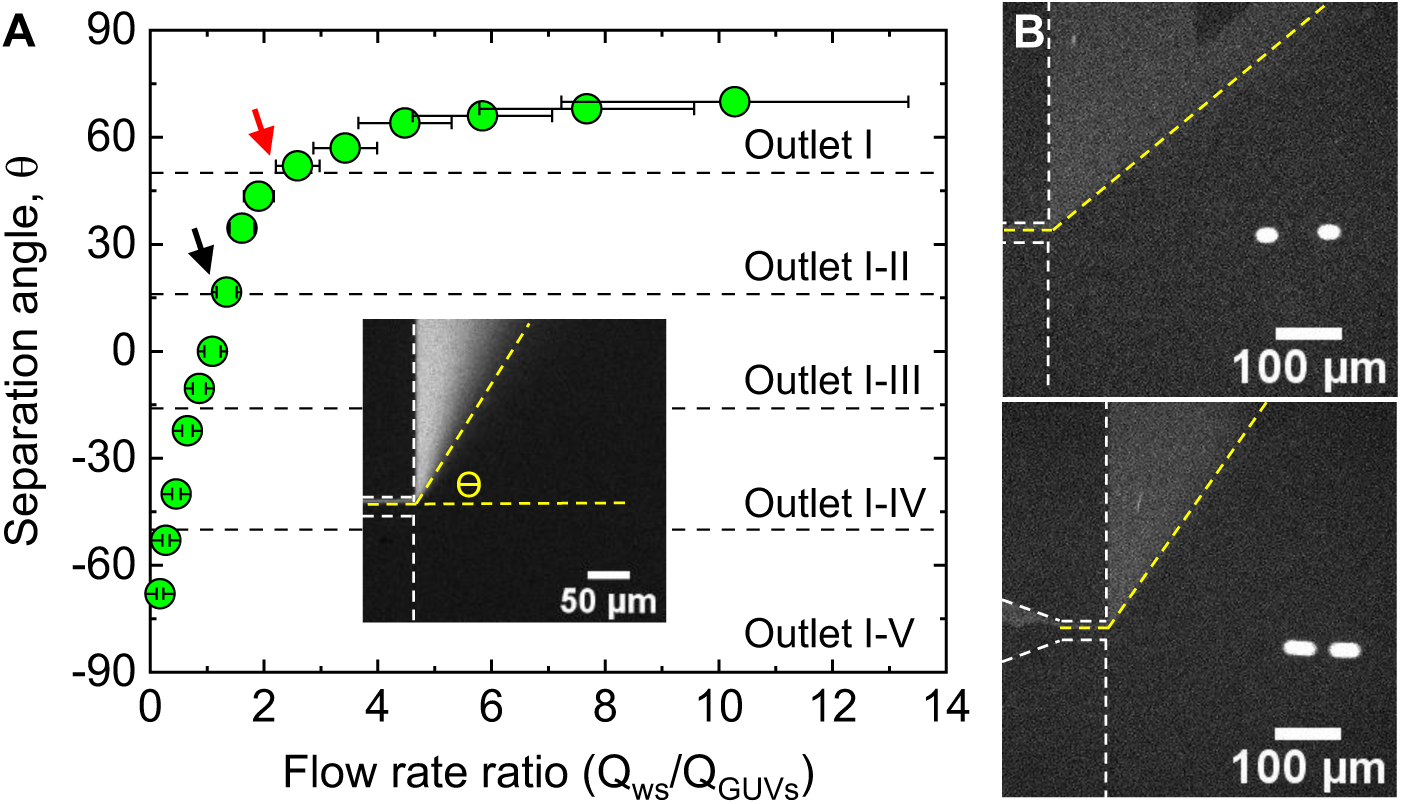
**A**. Measurements of separation angle *θ* as a function of flow rate ratio between the washing solution and GUVs mixture channels (*Q*_*ws*_/*Q*_*GUVs*_). The separation angle at different flow rate ratios was measured through focusing an HPTS solution using a second stream at variable flow velocities, as shown in the inset. The black and red arrows indicate the flow rate ratios at which the HPTS solution is bifurcated to outlet I&II and outlet I, respectively. **B**. Purification of GUVs at flow rate ratios of ∼1.6 (upper) and ∼2.6 (bottom), showing that the flow of GUVs to outlet III is not affected by the relative flow rate at the GUVs mixture and washing solution channels. The vesicles velocity *u* in each case, ∼0.0006 m s^-1^ (upper) and ∼0.001 m s^-1^ (bottom) was estimated from the figures using *u* = Δ*d*/Δ*t*, where Δ*d* is the vesicle displacement measured from the distance between the estimated centers of two circles (*a* = 23.3μm) that compose the blurred vesicle and Δ*t* is the camera exposure time which, in our setup, is inversely proportional to the frame rate (Δ*t =* 1⁄*FPS =* 0.025 *s*).

We next examined the mechanical stability of GUVs at the flow rate ratio range that allow effective vesicle separation (i.e., when *Q*_*ws*_⁄*Q*_*GUVs*_ > 1.3). Figure 4B shows the purification of GUVs at flow rate ratios of 1.6 and 2.6, where in both cases no vesicle breakage was observed after passing through the pinched segment. By taking *Q*_*GUVs*_ = 33 *μl* · *hr*^−1^ as a typical volumetric flow rate in the OLA-GUVs mixture channel (see section 3.2 and SI), the corresponding flow velocities in the pinched segment for each flow rate ratio are *v* = 0.039 *m s*^−1^ and *v* = 0.055 *m s*^−1^, respectively (SI). Using the appropriate capillary number 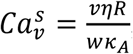, based on membrane stretching modulus *к*_*A*_, we can approximate the flow velocity at which the critical membrane tension of vesicles is reached (i.e., when rupture may occur). DOPC GUVs are known to retain their structural integrity at capillary numbers of 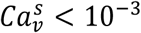 when exposed to viscous stress^*56, 57*^. By considering vesicles with *R* = 10μm and *к*_*A*_ = 0.25 N/m ^*55*^ in a pinched segment of *w* = 20μm, we obtain an upper bound of *v* ≈ 0.4 *m s*^−1^ for the flow velocity at which vesicle breakage may occur – far larger than the typical pinched segment flow velocities *∼*0.05 *m s*^−1^ (corresponding to *Q* = 100*μl/hr* in the pinched segment; see SI) used in the purification device. Hence, no vesicle breakage is expected to occur during continuous purification even when operated at large flow velocities of ca. 0.1 *m s*^−1^, which can be reached in the case of a very high vesicle production rate or when using the purification module as a standalone device (figure S1). Likewise, GUVs of other lipid compositions are also expected to remain stable under shear flow in our microfluidic platform^*58*^.

Still, under strong confinements (*2R* ≫ *w*) vesicles may experience much larger viscous forces and rupture as a result. We also note that while glycerol is known to increase the bending stiffness of lipid membranes^*59*^ and, thus, likely to enhance the stability of GUVs against rupturing, it is not a crucial component in the purification process and vesicles separation can be similarly achieved without it (figure S4 and section 3.5 below) as well as with different buffer compositions (SI).

### 3.5. Purification of polydisperse giant vesicles

Since vesicles separation in our purification module fundamentally relies on a narrow pinched-segment channel with dimensions comparable to the diameter of filtered GUVs, we examined whether it can be usefully applied to vesicle suspensions with a broad size distribution. To this end, we prepared a polydisperse giant vesicles suspension using electroformation, perfused it through a purification device (figure S1), and examined the filtered fraction with respect to purified OLA-GUVs with narrower size distribution.

Figure 5 shows representative confocal images and size distributions of OLA and electroformed vesicles before and after purification. As shown in figure 5A, nearly all octanol droplets were effectively removed from the OLA vesicles suspension, in accord with a purification efficiency of *e =* 0.99. The similar size distribution of OLA-GUVs before and after purification (figure 5B) implies that vesicles of different sizes – including vesicles with diameters smaller or larger than the pinched segment width (*a* ≈ *w* ± 5*μm*) – can be purified while maintaining their structural integrity. Likewise, we found that electroformed vesicles retain their stability when passed through the pinched-segment (*w* = 20*μm*) and could be separated from lipid aggregates that formed as a by-product during electroformation^*60*^ (figure 5C). Nevertheless, we note that in some cases a minute quantity of lipid aggregates or small vesicles (or any other type of residue) may not be excluded due to flow instabilities in the Y-junction or bursting of purified vesicles, as can be seen in figure 5C (after purification). Still, the amount of residual components that reach to outlet III can be diminished by closely monitoring the flow rate ratio and adjusting the flow rate of the focusing stream *Qws* accordingly.

**Figure 5.**
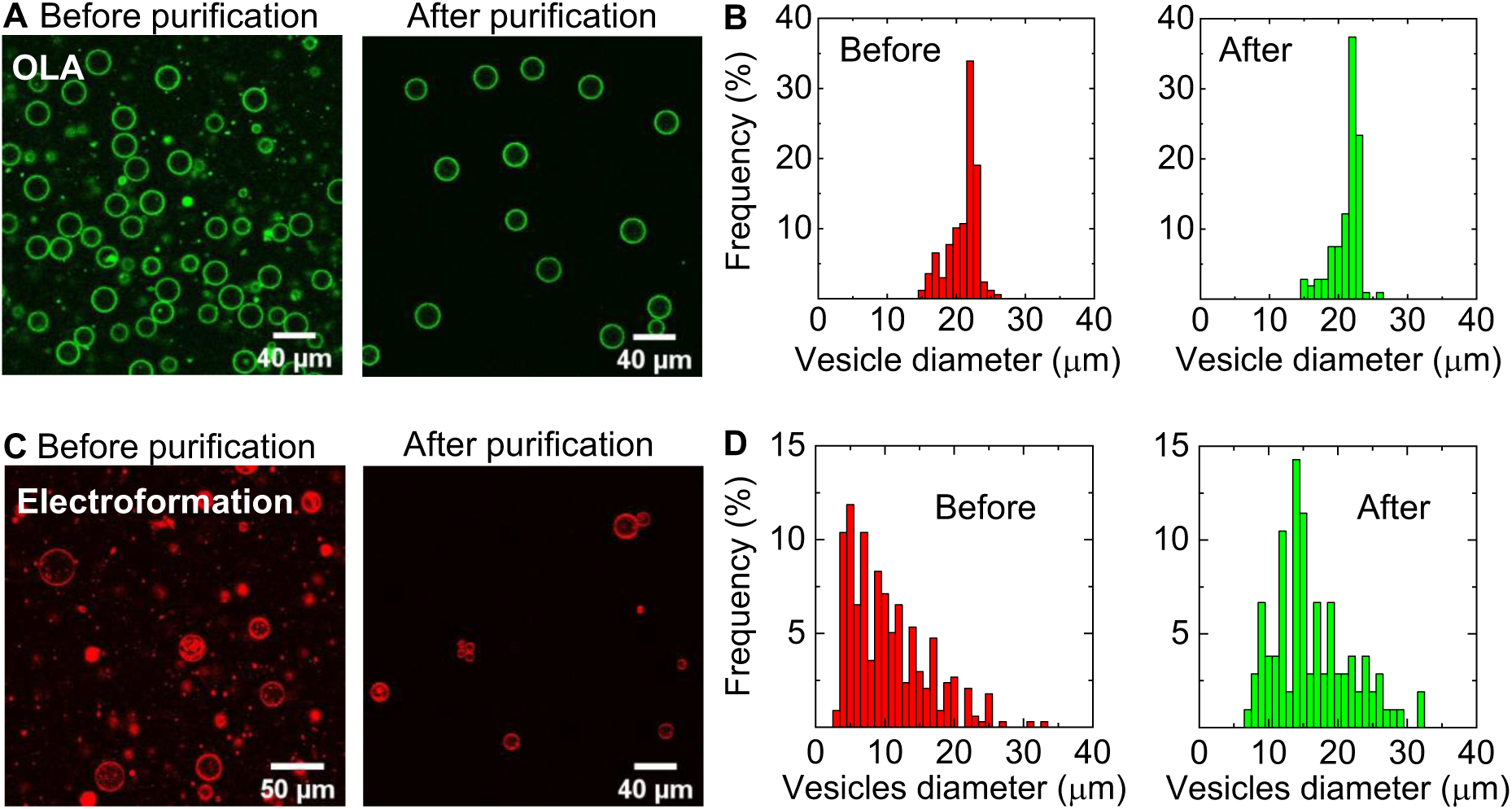
Separation of OLA-GUVs and electroformed-GUVs. **A**. Confocal images of OLA-GUVs before and after separation from octanol droplets. **B**. Frequency histogram of vesicle size distribution before (n=168) and after (n=107) purification. **C**. Confocal images of electroformed giant vesicles before and after purification showing the successful recovery of GUVs and exclusion of lipid aggregates. The washing solution was the same solution in which vesicles were prepared (experimental section). **D**. Frequency histogram vesicle size distribution before (n=337) and after (n=105) purification demonstrating the exclusion of giant vesicles with diameters *a* < 7*μm*. Assuming a gaussian distribution the average diameter of filtered vesicles is *ā* = 15 ± 9 *μm*.

A further examination of the electroformed vesicles size distribution before and after purification (figure 5D) indicates that vesicles with diameters smaller than 7*μm* were also efficiently removed from the mixture. In fact, some vesicles with diameters in the range 5*μm* ≤ *a* ≤ 20*μm* followed the spreading streamlines to outlet II; however, the fraction collected in outlet II also included lipid aggregates and therefore was not considered in the analysis (figure S5). In addition, we observed that vesicles or vesicle clumps with diameters much larger than the pinched-segment width got ruptured when crossing it (figure S5A), but importantly, no clogging was observed and chip operation was not obstructed as a result. Consequently, the purified vesicle fraction (outlet III) was obtained with a narrower size distribution and an average diameter of *a* ≅ 15 ± 9 *μm*, similar to OLA-GUVs, clearly indicating that on-chip purification is suitable for electroformed vesicles. However, as electroformed vesicles with diameters much smaller or larger than the pinched segment width (ca. *w* ± 0.5*w*) are either expelled or rupture during filtration, the final concentration of vesicles in the purified fraction is expected to be lower than in the untreated suspension. Conversely, since OLA-GUVs are produced with a narrow size distribution (figure 5B) their effective concentration can be largely retained after purification. Hence, while our purification technique can be utilized for polydisperse suspensions, it performs most efficiently when vesicles have a narrow size distribution with diameters comparable to the width of the pinched segment (i.e., *a* ≈ *w* ± 5*μm*).

### 3.6. Applicability of the integrated microfluidic device

Charge-mediated fusion between oppositely charged small unilamellar vesicles (SUVs) and GUVs is a contemporary technique for incorporating membrane proteins into the lipid bilayer of giant vesicles^*61, 62*^. Nevertheless, the presence of unfused SUVs may obstruct protein activity, adsorb (dock) to the vesicle membrane^*15*^, and introduce undesirable surface area that may interrupt with successive manipulation stages. We demonstrate the utility of our integrated microfluidic device by sequentially producing OLA-GUVs, fusing them with SUVs and then simultaneously purifying the fusion product from SUVs and octanol droplets.

Figure 6A shows the production of negatively charged OLA-GUVs (DOPC:DOPG 3:1 wt%, and 1 mol% NBD PE) and their subsequent fusion with positively charged SUVs (DOPE:DOTAP 4:1 wt%, and 0.1 mol% Liss Rhod PE) in the post-junction channel (figure 6A rightmost figure). We note that the full fusion between similar SUVs and OLA-GUVs as above was initially confirmed through internal content mixing measurement prior to mixing the vesicles in the integrated microfluidic chip (figure S6). Following their fusion on-chip, the vesicles reach the purification section (after ca. 5 min) where GUVs are simultaneously separated from SUVs and octanol droplets (figure 6B), demonstrating the usefulness of stream bifurcation for excluding different types of impurities regardless of their size. To quantify the fusion efficiency, we labelled the SUVs and GUVs membrane with the FRET pair DOPE-Rh (acceptor) and DOPE-NBD (donor), respectively. Upon fusion, DOPE-Rh is transferred from SUVs to the GUVs membrane, increasing the relative FRET efficiency *E*_*FRET*_ = *I*_*Rh*_/(*I*_*Rh*_ *+ I*_*NBD*_), where *I*_*Rh*_ and *I*_*NBD*_ are the fluorescence intensities of DOPE-Rh and DOPE-NBD following excitation of the latter. Figure 6C shows the incorporation of DOPE-Rh in the GUVs membrane after fusion in the integrated device. The transfer of Rh from SUVs to NBD labelled GUVs resulted in an increase of *E*_*FRET*_ from 0.25±0.02 to 0.*38*±0.09. Through measuring *E*_*FRET*_ at different DOPE-Rh concentration (figure S7), while maintaining the amount of DOPE-NBD in the GUVs membrane constant, we found that the fraction of DOPE-Rh transferred from SUVs to GUVs is 0.073mol%. By taking a GUV radius of *R* = 10 *μm* (see figure 5B), an SUV radius of *r* = 65*nm*, and assuming an area per lipid of 68 Å^*2*^ for all lipids, the estimated number of SUVs fused with a single GUV is *n(SUVs)* ≈ 1.7 × 104. likewise, it can be shown that for the same number of fused SUVs the number of positively charged lipids (DOTAP) transferred from SUVs (*n(DOTAP)* ≈ 3 × 10^8^ lipids) is comparable to the number of negatively charged lipids (DOPG) in GUVs ((*n(DOPG)* ≈ 4.5 × 10^8^ lipids)). Since DOPE-Rh transfer through full fusion is less likely to occur between positively charged SUVs and neutral GUVs^*15, 19, 63*^, the balance of negative charges in the fused GUVs membrane signifies that fusion time in the integrated device is sufficient for allowing efficient fusion between SUVs and GUVs.

**Figure 6.**
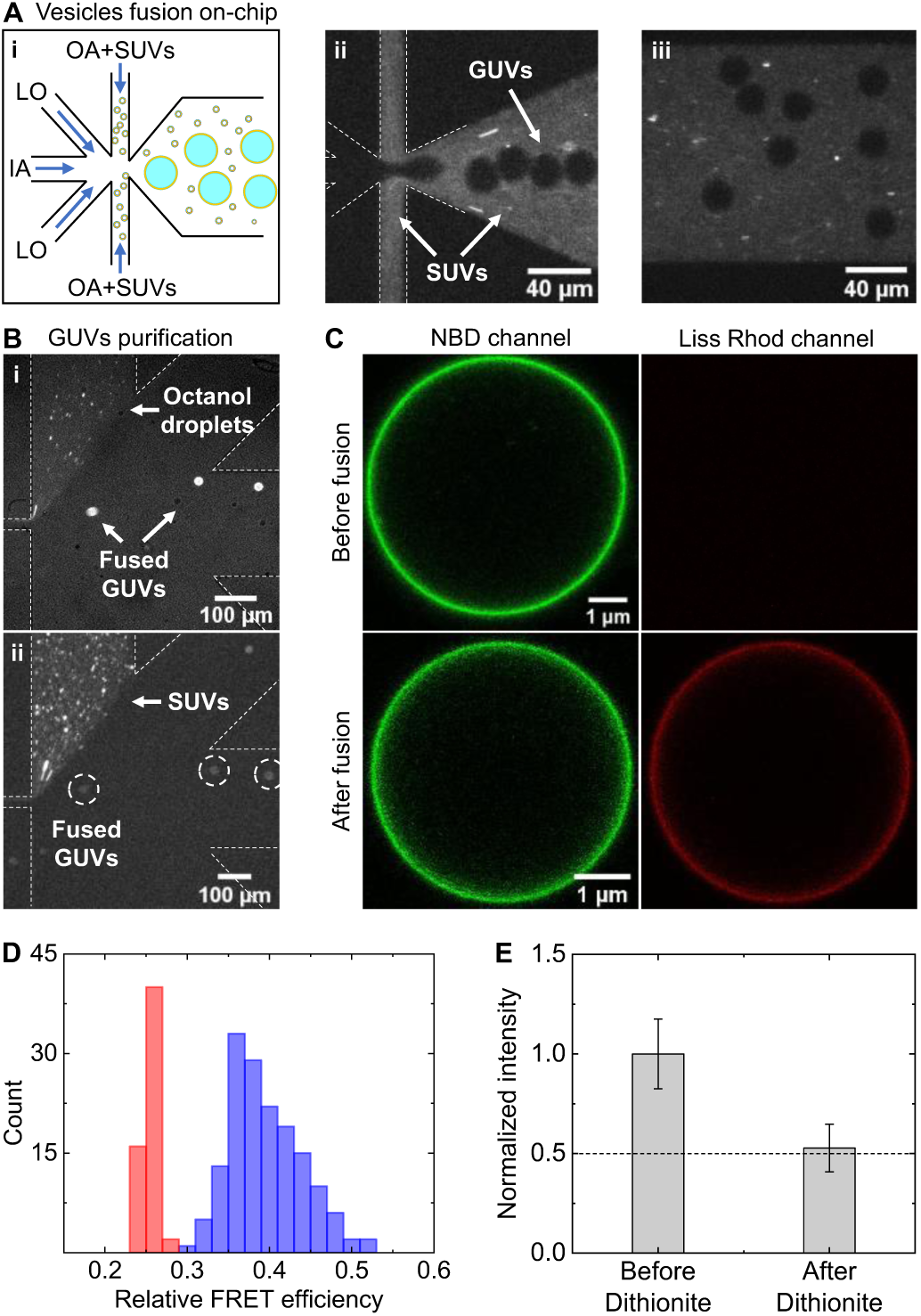
Charge-mediated vesicles fusion using the integrated production-purification microfluidic device. **A. (i)** A schematic showing the concept of on-chip fusion between positively charged SUVs and freshly-prepared negatively charged GUVs. **(ii)** A fluorescent image showing the production and fusion of DOPE-NBD labelled GUVs with DOPE-Rh labelled SUVs. SUVs are perfused through the outer aqueous channel. **iii**. A fluorescent image showing the mixing of SUVs and GUVs in the post-junction channel. Both images (ii and iii) were acquired at an excitation wavelength of 550nm to visualize the SUVs and the white dashed lines were added to illustrate the microchannels contour. **B**. Simultaneous separation of fused GUVs from octanol droplets and SUVs (*R* = 65 *nm*; zeta potential *ζ = +44 mV*) using the purification unit of the integrated device (figure S1). **C**. Confocal images (a magnified view) of GUVs, settled at the bottom of the imaging chamber, before (upper panel) and after (bottom panel) fusion with SUVs. The images were acquired at excitation wavelengths of 488nm and 559nm to illustrate the transfer of DOPE-Rh from SUVs to GUVs. **D**. Relative FRET efficiency (*E*_*FRET*_ = *I*_*Rh*_*/(I*_*Rh*_ + *I*_*NBD*_)) measurement before (red bars, n=57) and after (blue bars, n=192) fusion. The fluorescence intensity of Rh and NBD was measured after excitation of the later at wavelength of 488nm. **E**. Examination of membrane unilamellarity following on-chip fusion of positively charged SUVs (labelled with DOPE-NBD) and negatively charged GUVs, using dithionite reduction of NBD-PE lipids (see main text). The averaged fluorescence intensity of the fused GUVs membrane was measured before (n=28) and after 30 min (n=24) from the addition of dithionite. Fluorescence intensity was normalized based on the fluorescence intensity before the addition of dithionite.

To exclude the transfer of DOPE-Rh through hemifusion, we examined the degree of membrane unilamellarity using dithionite (S_2_O_4_^2-^), a membrane impermeable reducing agent, that reduces NBD and, as a result, renders it nonfluorescent. Following the addition of dithionite to unilamellar vesicles that contain NBD-labelled lipids, the fluorescence of the vesicles membrane is expected to decrease to 0.5 as only NBD in the outer leaflet is reduced. We produced negatively charged GUVs (DOPC:DOPG, 3:1) and fused them on-chip with NBD-labelled positively SUVs (DOPE:DOTAP, 4:1, with 1mol% DOPE-NBD). The fused GUVs were then extracted from the chip and the fluorescence intensity of DOPE-NBD was measured before and after the addition of dithionite. As can be seen in figure 6E, the average membrane fluorescence intensity was reduced to *0*.*52* ± *0*.*12* of its initial value, implying that SUVs and GUVs on-chip was governed by full fusion.

## 4. Conclusions

To conclude, we developed a PFF-based microfluidic platform capable of continuously purifying cell-sized vesicles through stream bifurcation, where one stream consists of all dissolved and dispersed extraneous components and the second contains giant vesicles in a residue-free solution. Fractionation of the bifurcated streams into five microchannel then allows the spatial separation and collection of purified vesicles. We showed that various residual components, from molecules to micron-size droplets, can be successfully removed with high efficiency (*e =* 0.99) based on their size and, in the case of oil droplets, also on their deformability. Notably, GUVs remain stable during the purification process and were not affected by the type of residues we examined. In addition, we demonstrated that the purification technique can be successfully applied to polydisperse suspensions of giant vesicles. By integrating the purification module with a microfluidic-based GUV-formation method we established a complete microfluidic unit that continuously produces and purifies GUVs. The relevance of our integrated device to synthetic biology is demonstrated through the sequential production, manipulation (SUVs fusion) and purification of GUVs from multiple residual components (oil droplets and unfused SUVs) at the same time. Altogether, our microfluidic-based purification method can be utilized either as a standalone device or as part of a microfluidic production-line for building artificial cell models from the bottom up.

## Acknowledgments

R. T. acknowledges funding from the European Union’s Horizon 2020 research and innovation programme under the Marie Sklodowska-Curie grant agreement No 892333, and from the Blavatnik Family Foundation. U. F. K. acknowledges support from an ERC consolidator grant (Designer-Pores 647144). K. A. N. acknowledges support from a Cambridge-National Physical Laboratory (UK) studentship, the Winton Programme for the Physics of Sustainability, the Trinity-Henry Barlow Scholarship, and the ERC. M. F. acknowledges support from the UK’s Engineering and Physical Sciences Research Council Doctoral Training Programme and a Cambridge-NPL case studentship. The authors thank Jack Webber for his help with profilometer measurements of our microfluidic designs.

## Author contribution

R.T. conceived the idea and designed the microfluidic devices and experiments. R.T. conducted the experiments and analyzed the data. M.F. conducted the chip simulations and bifurcation angle and flow rate ratio measurements. R.T. and M.F. conducted the vesicle fusion experiments. K.A.N prepared the electroformed GUVs. M.F., K.A.N and U.F.K assisted with theoretical aspects of the microfluidic purification system. R.T. wrote the manuscript. All authors discussed the results and commented on the manuscript.

## Electronic Supplementary Information (ESI) available

<u>S1:</u> Estimation of flow rates in the integrated device. <u>S2:</u> Separation of GUVs from oil droplets based on deformability. <u>S3:</u>Bifurcation angle and flow rate ratio measurements. <u>S4:</u> Effect of buffer composition on GUVs purification. <u>S5:</u> Purification of electroformed giant vesicles. <u>S6:</u> Internal content mixing after vesicles fusion. <u>S7:</u> Relative FRET efficiency calibration. <u>Figure S1:</u> Design of the standalone purification device. <u>Figure S2:</u> Flow rate measurement in the purification device. <u>Figure S3:</u> Separation of GUVs from large oil droplets. <u>Figure S4:</u> Sequential production and purification of GUVs in a buffer without glycerol. <u>Figure S5:</u> Purification of electroformed giant vesicles. <u>Figure S6:</u> Internal content mixing study following vesicles fusion. <u>Figure S7:</u>Relative FRET efficiency measurement.

