## Supplementary information for "A microfluidic platform for sequential assembly and separation of synthetic cell models"

### A microfluidic platform for continuous purification of cell-sized vesicles

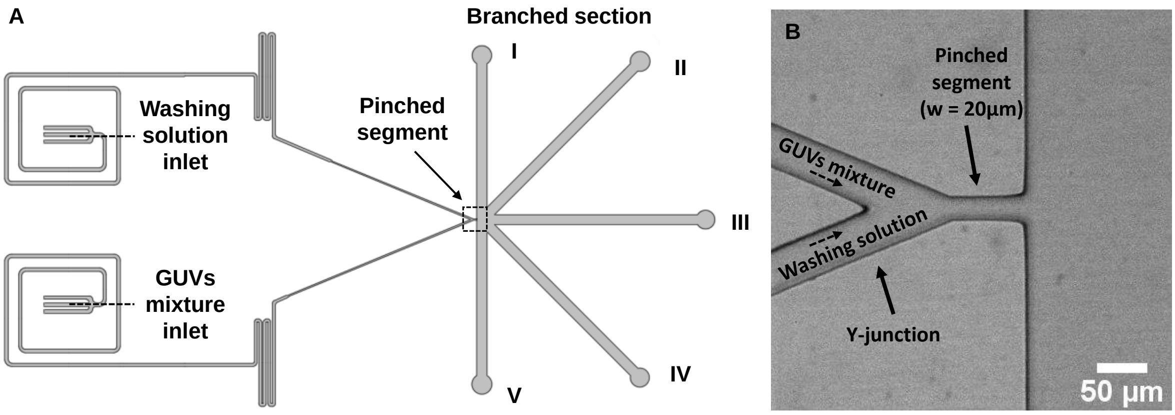


**Figure S1: A.** CAD design of the standalone purification device (using AutoCAD), with its principal components labelled. The device has 2 perfusion inlets through which the GUVs mixture and washing solution are perfused. The GUVs mixture is focused by the washing solution so vesicles flow to outlet III and impurities to outlets I and II (or just outlet I, depends on the flow rate ratio; see main text). **B.** A micrograph of the pinched segment (marked with a dotted line square in figure S1A) with the relevant components shown in the figure.

#### S1. Estimation of flow rates in the integrated device during GUVs production by OLA

To estimate the flow rates in the purification section of the integrated device (OLA-PFF) during GUVs production in the OLA section, we first evaluated the flow rate in the GUVs mixture channel $Q_{GUVs}$ by measuring the drift velocity of octanol droplets in the GUVs mixture channel (figure S2), as explained in the previous section. By measuring the bifurcation angle $\theta$ (see below), the flow rate ratio ($Q_{ws}/Q_{GUVs}$) at the pinched segment entrance can be evaluated (figure 4A). Subsequently, we can evaluate the flow rate in the washing solution channel $Q_{ws}$, for a given value of $Q_{GUVs}$, and the flow rate in the pinched segment using $Q_{ps} = Q_{ws} + Q_{GUVs}$ (for an incompressible fluid). Using the extracted flow rate values, we can then estimate the pressure at the entrance to the pinched segment through $P_{ps}^{in} = Q_{ps}(\mathrm{R_{ps}} + \mathrm{R_{bc}}) + P_{atm}$, where $\mathrm{R_{bc}}$ is the total hydraulic resistance of the five branched channels (figure 2A and figure S1) and $P_{atm}$ is the atmospheric pressure. Finally, by taking the value of applied pressure to the washing solution inlet, the pressure drop across each channel in the purification module can be estimated during a typical GUVs production by OLA. We found that during a typical GUVs production using OLA the mean flow velocity of the external fluid in the GUVs channel is $Q_{GUVs} = v_{drop} \times A_{GUVs} = 33.4 \pm 3.7\ \mu l\ hr^{-1}$, where $v_{drop} = 3.8 \times 10^{-3} \pm 4.2 \times 10^{-4}$ m $s^{-1}$ and $A_{GUVs} = 2400$ µ$m^2$. By taking a flow rate ratio of $Q_{ws}/Q_{GUVs} = 2$ (a typical value that allows to separate all impurities to outlet I; see figure 4A), the flow rate in the pinched segment during OLA can be estimated as $Q_{ps} = 3Q_{GUVs} = 100\ \mu l\ hr^{-1}$.

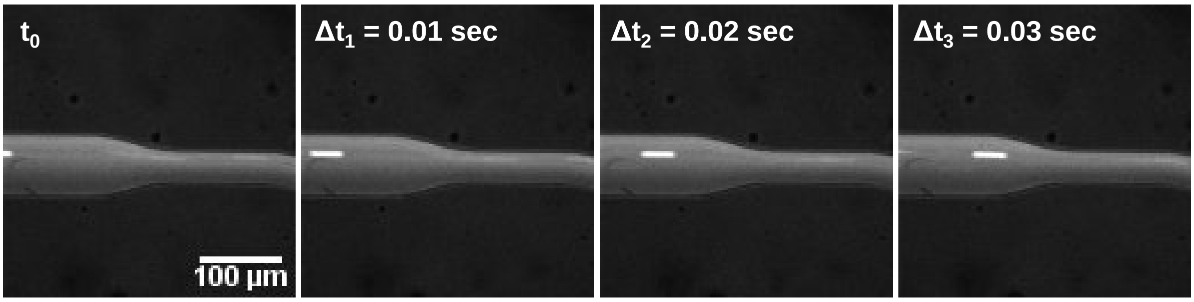


**Figure S2.** A time-lapse snap-shot of an oil droplet flowing in the GUVs mixture channel, where $\Delta t$ represent the time duration from $t_0 = 0$ sec. The oil droplet was generated at the OLA section and then drifted through the bridge channel to the purification module.

**S2. Separation of GUVs from oil droplets based on deformability**

The degree of droplet deformation can be viewed through the thickness $\delta$ of the uniform thin film that is formed between the drifting viscous drop and the pinched segment wall, where $\delta$ is determined by the capillary number that describes the balance between viscous stress and surface tension[1]. For octanol droplets flowing through the pinched segment, the capillary number is $Ca_d = \frac{v\eta_c}{\sigma} = 0.016$, where $\eta_c = 1.5 \times 10^{-3}\ pa \cdot s$ is the continuous phase viscosity (15% glycerol in water solution[2]), $v = 0.93\ m/s$ is the average velocity of the unperturbed external fluid (equivalent to a volumetric flow rate of $Q = 100\mu l/hr$ in the pinched segment (see previous sections), and $\sigma$ = 8.5mN/m is the interfacial tension between octanol and water[3] (this value is expected to be even lower, and thus $Ca_d$ higher, in the presence of the emulsifier poloxamer P188). Therefore, in the relevant regime $Ca_d^{-1/3} < \lambda < Ca_d^{-2/3}$, the film thickness between a confined octanol droplet with a radius $R = 0.5w = 10\mu m$ and one side of the pinched segment wall can be estimated as $\delta{\sim}1.3R(4Ca_d)^{2/3} = 2.2\mu m$.[4, 5] This suggests that large viscous octanol droplets with diameters comparable and even larger than the pinched segment width $(a/w \geq 1)$ elongate to occupy $1 - (2\delta/w){\sim}0.78$ of the microchannel width, as schematically illustrated in figure 1D. Consequently, their center of mass can be shifted by the focusing stream (washing solution) to the pinched segment sidewall – and away from the microchannel centerline – where the streamlines flow to outlets I and II (figure S3).

On the other hand, since vesicles are enclosed by an inextensible fluid membrane, they tend to elongate less along the flow direction and rather expand along the channel's less confined direction (i.e., the pinched segment height)[6, 7]. The large membrane stretching modulus[8, 9] retains the surface area and volume of vesicles nearly constant so their behaviour in Poiseuille flow is mainly dictated by their reduced volume $\mathcal{V} = V_{GUV}/V_{sphere}$ (a measure of vesicle asphericity) and the capillary number based on the membrane bending modulus $\kappa_b$, where $V_{GUV}$ and $V_{sphere}$ are the vesicle volume and the volume of a sphere with the same radius[7, 10]. Hence, for an undeformed GUV with a typical bending modulus[9] of $\kappa_b \approx 1 \times 10^{-19}$ J and radius of R=10μm in an average flow velocity $v = 0.93 m/s$ the capillary number is $Ca_v = \frac{v\eta R^4}{wh\kappa_b} = 15892$, where the viscosity is as above, w = 20 μm, and h = 42 μm. It was shown by Coupier et al.[7] that at similar capillary numbers confined vesicles ($a/w \geq 1$) in a rectangular channel, with a reduced volume of $\mathcal{V} \approx 1$ (i.e., nearly spherical immediately after crossing the pinched segment, as can be seen in figure S3 and figure 2B (iii)), may only slightly deform to favour a bullet shape geometry that retain their in-plane width roughly constant, contrary to droplets that adopt the aspect ratio of the channel. Therefore, since GUVs are expected to mainly extend along the pinched segment height (i.e., the less confined direction) with no significant reduction along the width direction, their center of mass should retain its position close to the microchannel centerline so they will drift to outlet III, as also shown in figure S3.

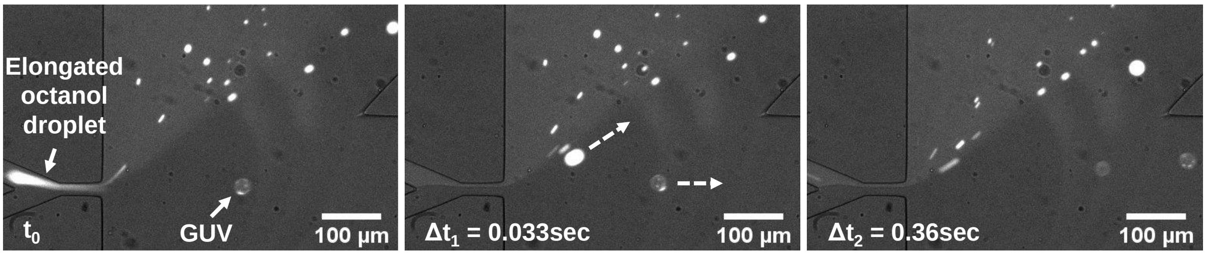


**Figure S3.** Purification of OLA-GUVs from oil droplets of different sizes, including dissolved octanol. **C.** A time-lapse snap-shot of large oil droplet ($a = 26\mu m$) separation from OLA-GUVs ($a = 24.5\mu m$), where $\Delta t$ represent the time duration from $t_0 = 0\ sec$.

**S3. Bifurcation angle and flow rate ratio measurements**

We evaluated the flow rate in the Y-junction channels and pinched segment by perfusing fluorescently tagged large unilamellar vesicles (LUVs; diameter: $a = 142 \pm 98\ nm$), as tracer particles, through both inlets of our purification device and measuring their drift velocity at different applied pressures. The GUVs mixture inlet channel was held at 1mbar and the washing solution inlet channel adjusted incrementally from 2mbar to 110mbar (for this measurement we used the purification unit of the integrated device where the GUVs mixture channel is shorter than the washing solution channel, and thus, its hydraulic resistance is much lower, allowing to achieve a fluid flow at a low applied pressure of 1 mbar). At each pressure step an image of the bifurcation angle $\Theta$ (see figure 4A) was taken as well as a short video (20 frames) of the flow close to each inlet were recorded. Flow velocities of individual particles were determined manually from the inlet flow videos by identifying particle displacements between a sequence of consecutive frames using ImageJ. The mean flow velocity $\langle v_c \rangle$ of several particles distributed evenly across the channel width approximated the mean velocity of the background fluid, where the subscript $c\ \in (ws, GUVs)$. By assuming an incompressible fluid, the flow rate in each inlet channel $Q_c$ as a function of applied pressure $P_{inlet}$ could be then calculated using $Q_c(P_{inlet}) = \langle v_c(P_{inlet}) \rangle\, A_c$, where $A_c$ is the cross-sectional area of the microchannel. Consequently, we could plot the bifurcation angles as a function of the flow rate ratio $(Q_{ws}/Q_{GUVs})$ between the two inlet channels (washing solution (ws) and GUVs mixture (GUVs)) as shown in Figure 4A. We note, however, that where the flow velocities of the tracer particles were too high to determine from the experimental videos, we exploited the linear relationships between the volumetric flow rate $Q_c$ and the variable pressure applied to the washing solution channel $P_{ws}$ to extrapolate to high values of $P_{ws}$.

#### S4. Effect of buffer composition on GUVs purification

We used a broad range of buffer compositions (with and without glycerol) to examine the effect of the buffer on the stability of GUVs during their preparation and separation from residues. In all the following buffers stable GUVs production and purification were obtained (data is shown in the main text only for the first two compositions): 10mM HEPES (with and without glycerol); 100mM sucrose solution in miliQ; 10mM Tris, 150mM NaCl (with glycerol); 10mM Tris, 1mM EDTA (without glycerol); 10mM Tris, 50mM LiCl (without glycerol); 10mM Tris, 50mM LiCl, 2.5mM $MgCl_2$ (without glycerol); and 10mM Tris, 150 LiCl (without glycerol). Figure S4, shows the purification of DOPC:DOPG GUVs in 10mM HEPES (pH = 7.6), 200mM sucrose (without glycerol).

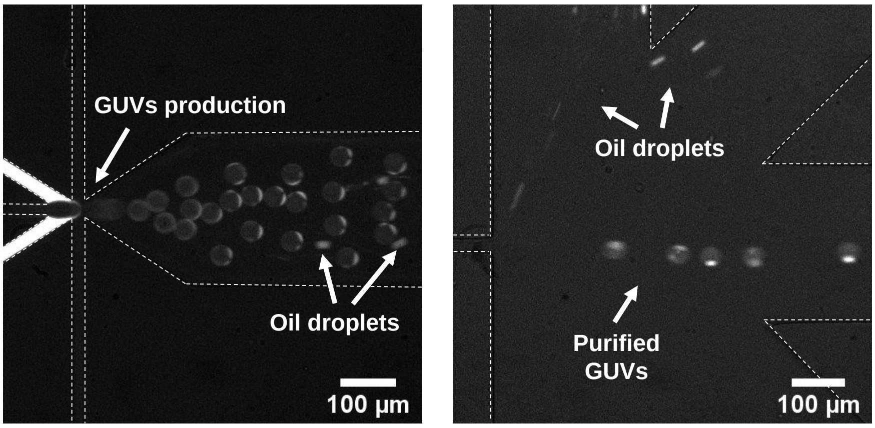


**Figure S4.** Sequential production and purification of DOPC:DOPG (3:1wt%) GUVs in a buffer without glycerol, using the integrated device shown in figure 2 (main text). The GUVs membrane was fluorescently labelled with 18:1 Liss Rhod PE and micrographs were acquired using an epifluorescence microscope.

#### S5. Purification of electroformed giant vesicles

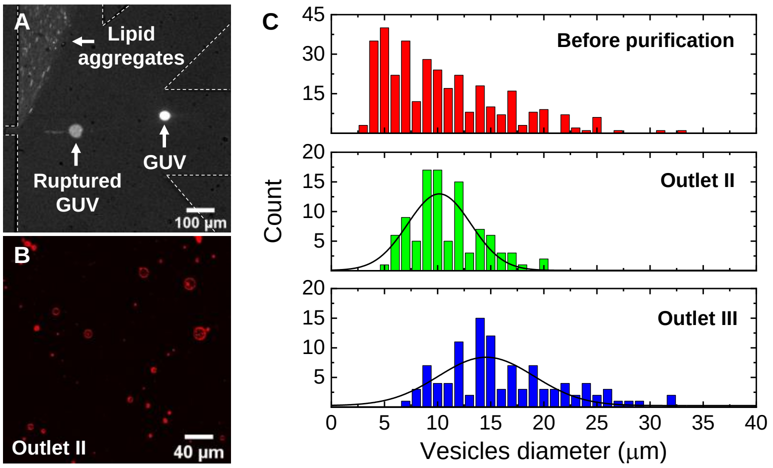


**Figure S5. A.** On-chip separation of Liss Rhod PE labelled electroformed giant vesicles from lipid aggregates using a standalone purification module. GUVs larger than ca. $w + 0.5w$ (i.e., ca. a >30 µm) were ruptured once crossing the pinched segment ($w = 20$ µm). **B.** Confocal images of the electroformed giant vesicles (GVs) and lipid aggregates which extracted from outlet II. **C.** Vesicle size distribution before (n=337) and after purification (Outlet II: n=100; outlet III: n=105). Assuming a gaussian distribution, the average diameter of filtered vesicles in outlet II is $\bar{a} = 10.2 \pm 5.9\ \mu m$ and in outlet III is $\bar{a} = 14.6 \pm 9.0\ \mu m$.

#### S6. Internal content mixing after vesicles fusion

Fully fused vesicles merge their internal content to form a larger interconnected vesicle with a unilamellar membrane. We verified the full fusion between negatively charged GUVs (DOPC:DOPG 3:1) and positively charged SUVs (DOTAP:DOPE 1:4) through measuring their internal content mixing using the calcein-cobalt-EDTA method[11]. When $Co^{2+}$ ions are complexed with the fluorophore calcein its fluorescence dramatically quenches. However, in the presence of EDTA, which preferentially chelates with $Co^{2+}$, the fluorescence intensity of calcein increases[11, 12]. Hence, in the case of full fusion between EDTA loaded negatively charged GUVs and positively charged SUVs loaded with the $Co^{2+}$-calcein, we expect to observe a significant increase in the fluorescent intensity of fusion product's lumen compared to before fusion, as schematically depicted in figure S5A. To this end, we prepared GUVs loaded with 10mM EDTA in a PBS buffer (10mM PBS, 15% glycerol, 200mM sucrose) and

SUVs loaded with 1mM $Co^{2+}$-calcein in miliQ water. The GUVs were mixed with SUVs in an isosmotic glucose buffer (10mM PBS, 15% glycerol, 200mM glucose) and were left to sediment to the bottom of an observation chamber by the density gradient between glucose (external solution) and sucrose (GUVs internal solution). As can be seen in figure S5B, the lumen of the fusion product became strongly fluorescent (figure S5B(ii)) upon fusion, compared to the GUVs lumen before mixing with SUVs (figure S5B(i)). Furthermore, we verified that such an increase is not due to permeation of the free calcein-cobalt complex through the GUVs membrane (figure S5B(iii)), indicating that full fusion indeed occurs between SUVs and GUVs.

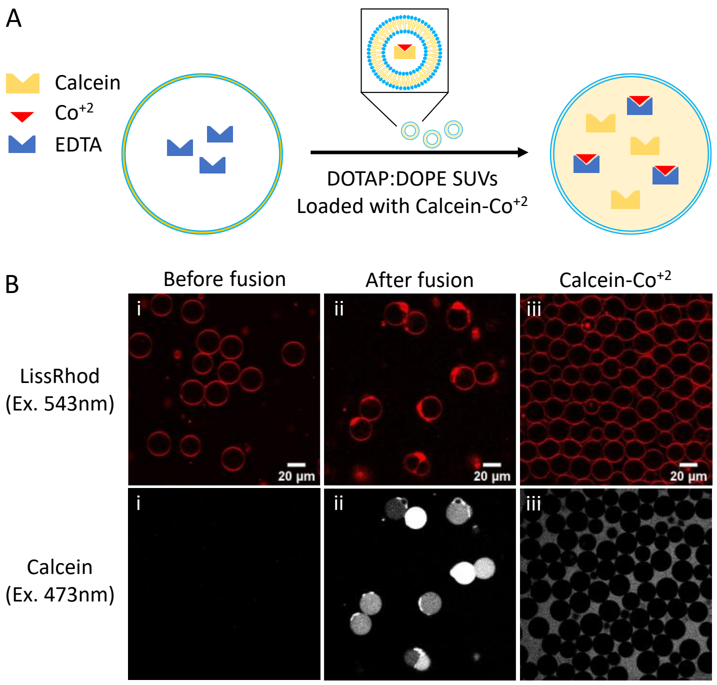


**Figure S6.** Internal content mixing study following vesicles fusion in an incubation chamber. **A.** Schematic illustration of the calcein-cobalt assay for studying internal content mixing. When $Co^{2+}$ ions are complexed with the fluorophore calcein its fluorescence is dramatically quenched; however, in the presence of EDTA, which preferentially chelates with $Co^{2+}$, calcein fluorescence is regained. **B.** Internal content mixing measurement after fusion between EDTA loaded GUVs and SUVs loaded with the $Co^{2+}$-calcein complex. The appearance of a fluorescent signal in the lumen of GUVs can be clearly seen after fusion with SUVs (ii, bottom panel), while no fluorescence can be detected in the vesicles lumen either before fusion (i) of after adding the $Co^{2+}$-calcein complex to the external solution of GUVs (iii).

#### S7. Relative FRET efficiency calibration

To correlate the membrane concentration of DOPE-Rh to the relative FRET efficiency, we prepared negatively charged OLA-GUVs (DOPC:DOPG, 3:1) with 1mol% DOPE-NBD and different concentrations of DOPE-Rh that range between 0 to 0.5 mol%. The relative FRET efficiency $E_{FRET} = I_{Rh}/(I_{Rh} + I_{NBD})$ was then calculated by measuring the DOPE-NBD and DOPE-Rh fluorescence intensities ($I_{NBD}$ and $I_{Rh}$, respectively) following excitation of NBD with a 488nm laser.

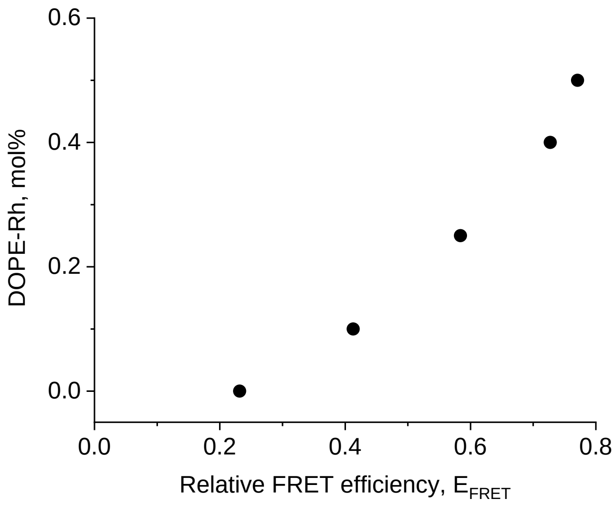


**Figure S7.** Relationship between DOPE-Rh concentration in GUVs membrane (contains 1mol% DOPE-NBD) and relative FRET efficiency.
